# Cold Shock Domain Protein LIN-66 cooperates with microRNA-pathway buffering to safeguard developmental timing

**DOI:** 10.64898/2026.07.07.737059

**Authors:** Reyyan Bulut, Victor Ambros

## Abstract

Robust execution of developmental cell fates requires precise spatiotemporal control of the fate-defining regulators. In *Caenorhabditis elegans*, temporal patterning of larval hypodermal fates is governed by the heterochronic gene regulatory network, in which microRNAs act as major post-transcriptional regulators by silencing temporal transcripts through 3**′**UTR-dependent repression. Here, we investigate *lin-66*, which encodes a nematode-specific cold shock domain protein previously implicated in heterochronic regulation and reported to associate with the miRISC effector protein AIN-1. Using targeted domain mutations and genetic analysis, we show that LIN-66 activity in the hypodermal cell-fate patterning requires its cold shock domain. Loss of *lin-66* causes persistent expression of LIN-14 and LIN-28, two early temporal regulators in the hypodermal seam cells that are canonical microRNA targets. Analysis indicates that *lin-66* function in seam-cell fate patterning does not depend on the native 3**′**UTR sequences of *lin-14* or *lin-28*, distinguishing its activity from canonical microRNA repression.. Consistent However, consistent with the a broad functional overlap between LIN-66 function and microRNA-mediated regulation, hypodermal *lin-66* loss-of-function phenotypes are strongly enhanced by mutations in *alg-1* and *ain-1/2*, which encode components of the microRNA-induced silencing complex. Moreover, loss of *lin-66* enhances phenotypes in mutants sensitized for microRNA activity outside the hypodermis. Together, these findings identify LIN-66 as a cold shock domain-dependent post-transcriptional regulator that safeguards developmental timing by limiting persistence of early fate regulators through mechanisms that intersect with, but are partly separable from, canonical 3**′**UTR-mediated microRNA repression.

## INTRODUCTION

Cold shock domain proteins can couple RNA binding to diverse post-transcriptional regulatory outputs, including mRNA stability, localization, and translation (Heinemann & Roske, 2021). LIN-66 has emerged as one such regulator in *Caenorhabditis elegans*. LIN-66 is a nematode-specific protein with extensive intrinsically disordered regions (IDRs) and a predicted cold shock domain, previously implicated in both developmental timing and protein translation in motor neurons (Blazie et al., 2024a; Morita & Han, 2006). In the larval hypodermis, loss of *lin-66* disrupts the temporal patterning of vulval and seam cell divisions, phenotypes previously attributed to de-repression of the heterochronic RNA-binding protein LIN-28 (Morita & Han, 2006). In cholinergic motor neurons, however, LIN-66 appears to act in the opposite direction, promoting translation of a subset of EIF-3.G-dependent mRNAs (Blazie et al., 2021, 2024b). Structure-function analyses further identified the cold shock domain as a critical functional module: mutations in conserved residues within the β2 strand and β2–β3 loop, which include the RNP1 RNA-recognition surface (Heinemann & Roske, 2021; Landsman, 1992), abolish neuronal LIN-66 function, whereas deletion of the flanking IDRs alters protein abundance and localization but preserves activity in that context (Blazie et al., 2024b). Together, these findings suggest that LIN-66 acts by binding to RNA to regulate gene expression, but how this activity contributes to temporal cell-fate control remains poorly understood.

At hatching, *C. elegans* larvae contain 10 seam cells on each lateral side; these cells undergo a stereotyped series of asymmetric and symmetric divisions during larval development. At the second larval stage (L2), the V1–V4 and V6 seam cells divide symmetrically, increasing the total number of seam cells to 16 per side (Sulston & Horvitz, 1977). The heterochronic gene governs temporal patterning of seam cell fates (Ambros & Horvitz, 1984), in part through microRNAs that post-transcriptionally downregulate key transcription factors and other temporal regulators (Abbott et al., 2005; Lee et al., 1993; Reinhart et al., 2000; Wightman et al., 1993). In *C. elegans*, microRNA-mediated repression is executed by the microRNA-induced silencing complex (miRISC), whose core components include microRNA-bound Argonaute (AGO) proteins ALG-1 and ALG-2, and GW182-like adaptor proteins AIN-1 and AIN-2. miRISC function is further supported by associated cofactors and effector complexes, including Poly(A)-binding protein (PABP), the CCR4-NOT deadenylase complex (Bartel, 2018), the TRIM-NHL family protein NHL-2 (Hammell et al., 2009), and GYF-domain protein GYF-1 (Mayya et al., 2021).

The heterochronic transcription factor HBL-1 promotes the L2-specific symmetric seam cell divisions (Abrahante, Daul, Li, Volk, Tennessen, Miller, Rougvie, et al., 2003; Lin et al., 2003). HBL-1 is developmentally downregulated through multiple regulatory inputs, including post-transcriptional repression by *lin-4* and *let-7*-family microRNAs (Abbott et al., 2005; Karp & Ambros, 2012; Lin et al., 2003), and post-translational regulation by LIN-46 (Ilbay & Ambros, 2019b; Pepper et al., 2004). These HBL-1 regulatory mechanisms are themselves embedded in a feedback-rich network in which LIN-28 represses *let-7*-family microRNA expression (Nelson & Ambros, 2019; Stefani et al., 2015; Van Wynsberghe et al., 2011) and LIN-46 (Ilbay et al., 2021). LIN-28, LIN-14, and *let-7-family* microRNAs also form an interconnected feedback module that reinforces early developmental states: LIN-14 negatively regulates *let-7-family* microRNAs, *let-7-family* microRNAs post-transcriptionally repress *lin-28*, and LIN-28 promotes LIN-14 expression, perhaps through direct binding to *lin-14* mRNA (Arasu et al., 1991; Stefani et al., 2015; Tsialikas et al., 2017). Both LIN-28 and LIN-14 are also post-transcriptionally repressed by *lin-4* after the L1 stage (Arasu et al., 1991; Lee et al., 1993; Moss et al., 1997; Wightman et al., 1993). Previous work placed *lin-66* upstream of *lin-28* in the heterochronic pathway. In *lin-66(lf)* mutants, LIN-28 persists ectopically after the L2 stage without a corresponding increase in *lin-28* mRNA abundance, and reporter analyses suggested that *lin-66* represses *lin-28* through the *lin-28* 3’UTR (Morita & Han, 2006). LIN-66 has also been recovered in association with proteins that function in or alongside miRISC, including AIN-1, NHL-2 and the scaffolding subunit of the CCR4-NOT deadenylase complex NTL-1 (Mayya et al., 2021; Wu et al., 2016). These observations suggested that LIN-66 may associate with microRNA-regulated mRNAs *in vivo*.

Here, we show that LIN-66 requires its cold shock domain to promote proper temporal patterning of hypodermal seam cell fates. Loss of *lin-66* causes inappropriate persistence of the early temporal regulators LIN-28 and LIN-14, and genetic epistasis reveals that *lin-66* has a major effect on developmental timing through regulation of *lin-14*. We further show that *lin-66* genetically interacts with multiple components of the microRNA pathway and enhances microRNA-sensitive phenotypes in both hypodermal and non-hypodermal developmental contexts. However, *lin-66* function in the seam cell fate patterning does not require the native 3’UTRs of *lin-28* or *lin-14*, indicating a relatively indirect interaction of LIN-66 with canonical microRNA repression. Together, these findings identify LIN-66 as a cold-shock-domain-dependent post-transcriptional regulator that acts in functional overlap with miRISC-associated regulation to reinforce developmentally timed gene expression dynamics underlying cell fate timing programs.

## RESULTS

### The function of LIN-66 in the temporal regulation of seam cell fates requires the cold shock domain

A BLAST search of the LIN-66 primary amino acid sequence against available protein databases did not identify predicted sequence homologs outside of the nematode phylum. Structural prediction of LIN-66 using the Rosetta web server (Kim et al., 2004), followed by structural alignment to the Protein Data Bank (PDB) using the DALI server (Holm et al., 2023), confirmed previous findings that LIN-66 contains a cold shock domain (CSD) (Blazie et al., 2024) (Fig. 1A). To examine conservation of this domain, we aligned LIN-66 CSD sequences from 49 *Caenorhabditis* species, revealing extensive conservation across the domain (Fig. 1B). We then compared the *C. elegans* LIN-66 CSD with CSDs from well-characterized CSD proteins, including human LIN-28A, *C. elegans* LIN-28A, *Xenopus tropicalis* LIN-28B, human YBX1, and *Bacillus subtilis* Csp1. This comparison showed that the hallmark residues conserved among canonical CSD proteins are also conserved in *C. elegans* LIN-66. The LIN-66 CSD shows additional conservation across *Caenorhabditis* species beyond these core CSD residues (Fig. 1C).

**Figure 1:**
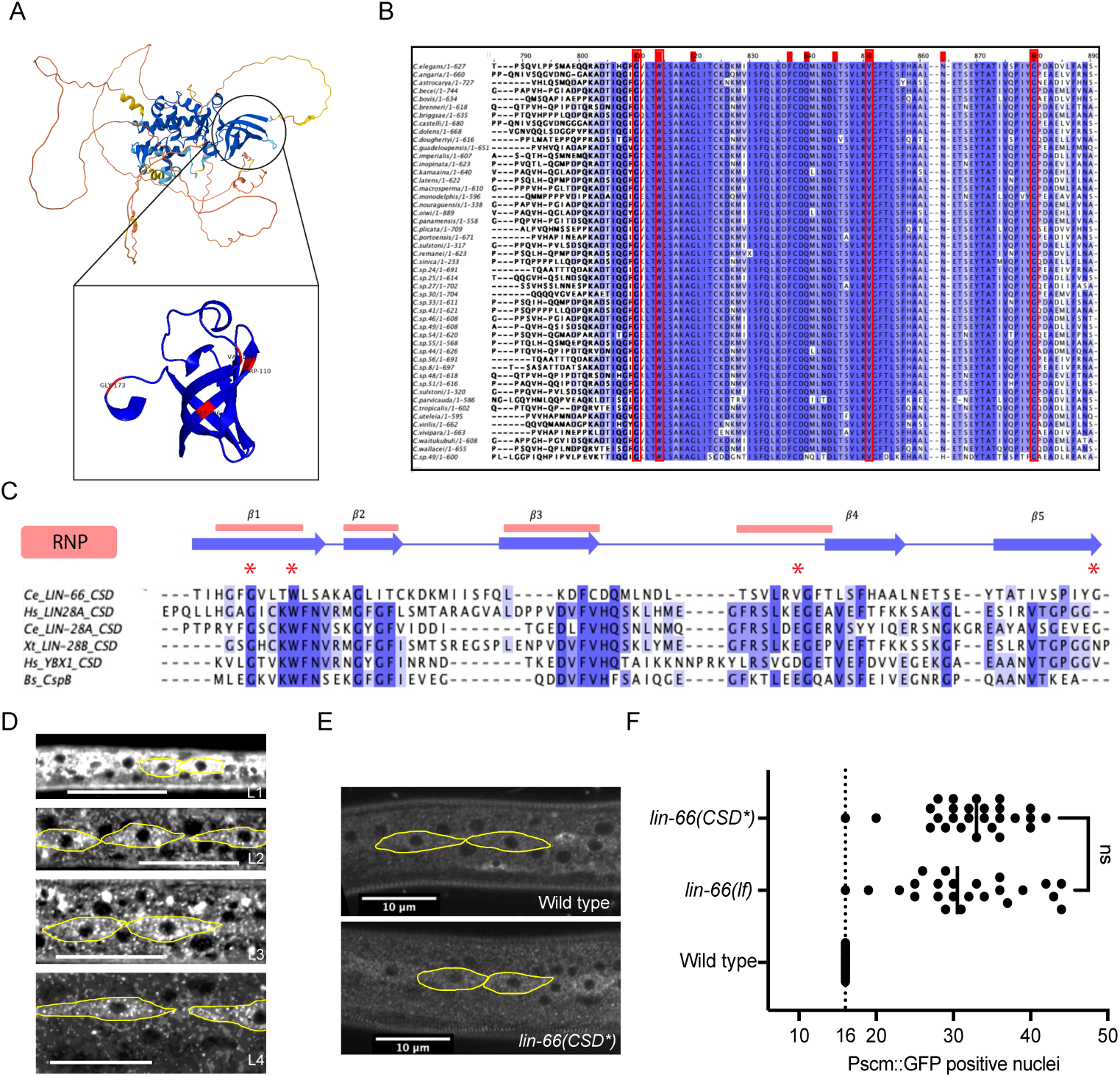
LIN-66 Cold Shock Domain is deeply conserved among *Caenorhabditis* species and required for the *lin-66* seam cell fate regulatory function. **A:** Putative three-dimensional structure of LIN-66 involves extensive disordered regions and a cold shock domain fold. **B:** CSD residues are deeply conserved among 49 *Caenorhabditis* species. **C:** Multiple sequence alignment and secondary structure annotation of cold-shock domains of *C. elegans* LIN-66 and LIN-28, human Lin28 and YBX1, *Xenopus tropicalis* LIN-28, and *Bacillus subtilis* CspB. Red asterisk indicates the residues that are mutated to alanine in *lin-66(CSD*)*. **D:** The seam cell expression of LIN-66::GFP is visible in all larval stages and displays a puncta-like pattern. **E:** LIN-66::GFP expression pattern is not changed in the cold shock domain mutant in comparison to wild-type worms. **F:** Number of *Pscm::GFP* positive nuclei in wild type, *lin-66(ok3326),* and *lin-66(CSD*)* mutant worms. *lin-66(CSD*)* phenocopies *lin-66(lf)* in seam cell overproliferation phenotype.

To examine LIN-66 expression and subcellular localization, we generated an endogenous *lin-66::GFP* that preserves the native *lin-66* regulatory sequences employing a flexible (Ser-Gly-Gly-Gly)₃ linker between LIN-66 and the c-terminal GFP tag. Animals carrying this allele were phenotypically indistinguishable from wild type, indicating that the tagged protein retains LIN-66 function. LIN-66::GFP was broadly expressed throughout development, with visible signal in the hypodermis, germline, and neurons (data not shown), and appeared predominantly cytoplasmic. In seam cells, LIN-66::GFP signal was most prominent during L1 and remained detectable at later larval stages, where it appeared broadly similar in abundance (Fig. 1D).

To test the functional requirement for the cold shock domain, we substituted four conserved residues within the predicted CSD —Gly106, Trp110, Val147, and Gly173— with alanine in the endogenous GFP-tagged *lin-66* locus, generating *lin-66(CSD*)* (Figs. 1A, C, E & F). LIN-66(CSD*) retained a localization pattern similar to wild-type LIN-66, appearing diffusely distributed in the cytoplasm and enriched in cytoplasmic puncta (Fig.1 E). Despite this apparently normal localization, *lin-66(CSD*)* animals exhibited excessive seam cell proliferation comparable to that of the *lin-66(lf)* mutant (Fig. 1F). Thus, these conserved CSD residues are required for LIN-66 function in temporal seam cell fate patterning.

### *lin-66(lf)* results in temporal dysregulation of LIN-28::GFP and LIN-14::GFP

To determine whether *lin-66* affects the expression of key regulators of early larval seam cell fate transitions, we examined endogenously tagged LIN-28::GFP, LIN-14::GFP and HBL-1::mScarlet-I in wild-type and *lin-66(lf)* larvae. Previous western blot analysis showed ectopic LIN-28 protein expression in *lin-66(lf)* mutants without a corresponding increase in lin-28 mRNA abundance, suggesting that LIN-66 limits LIN-28 accumulation post-transcriptionally (Morita & Han, 2006). Consistent with this prior observation, endogenously tagged LIN-28::GFP was elevated in the seam cell cytoplasm of *lin-66(lf)* animals at the L3 stage (Fig. 2A).

**Figure 2:**
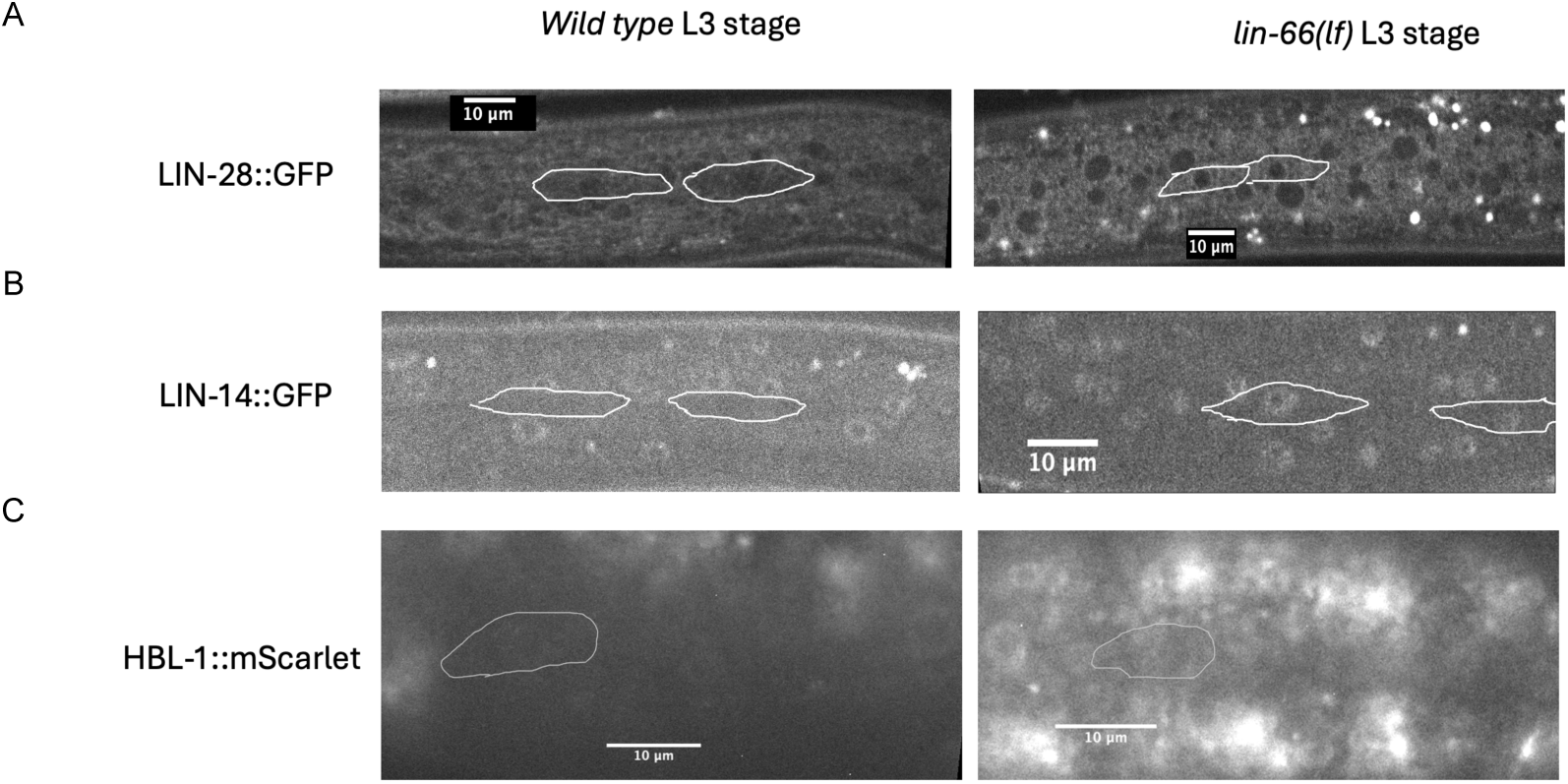
L3 larval stage expression patterns of early temporal regulators in wild-type and *lin-66(lf)* animals. Example seam cell borders are circled/marked in gray. **A:** LIN-28::GFP is modestly upregulated in seam cell cytoplasm at the L3 stage in *lin-66(If*) mutants. **B:** LIN-14 is not visible in the seam cells after the L2 stage in wild-type animals. *lin-66(lf)* animals ectopically express LIN-14::GFP in seam cells at L3 stage. **C:** HBL-1::mScarletI is down-regulated in the seam cells at the L3 stage in both wild-type and *lin-66(If*) mutants.

The effect of *lin-66(lf)* on LIN-14 expression was more pronounced than for LIN-28. Whereas LIN-14::GFP was not detected in wild-type seam cells at the L3 stage, *lin-66(lf)* animals showed ectopic LIN-14::GFP expression (Fig. 2B) and beyond, indicating that *lin-66* is required for the timely downregulation of LIN-14 during larval development. By contrast, we did not detect a difference in HBL-1::mScarlet-I expression between wild-type and *lin-66(lf)* animals (Fig. 2C). Thus, among the early temporal regulators examined, loss of *lin-66* most predominantly disrupts LIN-14 downregulation, with a more modest effect on LIN-28 and no detectable effect on HBL-1 expression.

### Perturbations of the heterochronic gene cascade in *lin-66(lf)* mutants

Previous genetic analysis placed *lin-66* upstream of *lin-28* in seam cell fate regulation (Morita & Han, 2006). However, the heterochronic gene network known to control seam cell patterning has expanded (Fig. 3A) to include an interconnected *lin-14-let-7-family-lin-28* feedback module and multiple regulatory arms controlling HBL-1 (Ilbay et al., 2021; Ilbay & Ambros, 2019b; Tsialikas et al., 2017). We therefore revisited the genetic position of *lin-66* within this expanded network.

**Figure 3:**
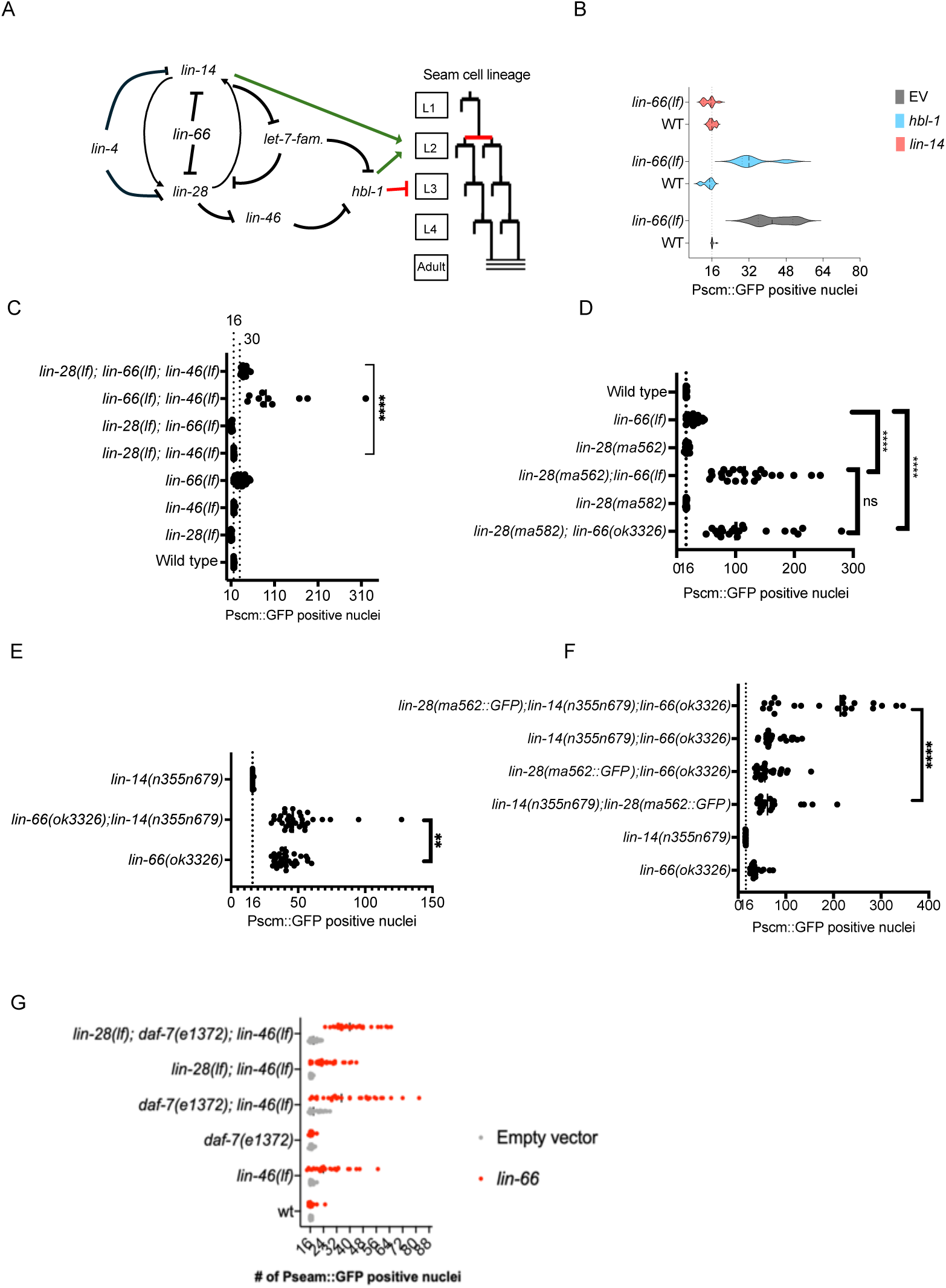
Perturbations of the heterochronic gene cascade in *lin-66(lf)* mutants. **A:** Genetic model for regulatory relationships among genes governing seam cell lineage fate in wild-type *C. elegans* hermaphrodites based on previous publications and the results of experiments reported here. Seam cells divide asymmetrically at the beginning of each larval stage, with a L2-specific symmetric division occurring just before the asymmetric division in the L2. The L2-specific seam cell symmetric division is specified by dosage-dependent activities of *lin-14* and *hbl-1*. According to the model, *lin-14* and *hbl-1* exert negative regulation of transitions from the L1 seam cell fate (no symmetric division) to the L2 seam cell fate (symmetric division), and from the L2 (symmetric division) to the L3 seam cell fate (asymmetric division). Previous genetic analysis of *lin-14* mutants suggests that transition from L1 to L2 fates requires downregulation of *lin-14* from highest level to the reduced but still active level (Ambros & Horvitz, 1987). Transition from L2 to L3 fates requires full downregulation of LIN-14 and HBL-1 (Abrahante, Daul, Li, Volk, Tennessen, Miller, & Rougvie, 2003; Ilbay & Ambros, 2019b; Seggerson et al., 2002). *lin-28* plays a central role in downregulation of LIN-14 and HBL-1, by promoting LIN-14 expression, likely by binding to the *lin-14* transcript (Stefani et al., 2015), and supports HBL-1 expression via a circuit through the post-translational regulator, LIN-46 (Ilbay & Ambros, 2019b). Developmentally expressed microRNAs, particularly *lin-4* and the *let-7 family* (which are upregulated in the L1 and L2 stages, respectively (Tsialikas et al., 2017), trigger the developmental downregulation of LIN-14, LIN-28, and HBL-1. Our data suggests that *lin-66* wild-type activity contributes to controlling the timing of the L2 to L3 fate transition by post-transcriptional negative regulation of *lin-14* and *lin-28* expression. **B:** Seam cell phenotypes of *lin-66(lf)* following of *lin-14* and *hbl-1* RNAi. *lin-14* RNAi suppresses the extra seam cell phenotypes of *lin-66(lf)* animals, whereas *hbl-1* does not. **C:** *lin-66(lf)* significantly aggravates the seam cell defects of both the *lin-46(lf)* single mutant and the *lin-28(lf); lin-46(lf)* double mutant. *lin-28(lf); lin-66(lf)* animals phenocopy *lin-28(lf).* **D:** Loss of *lin-66* results in excessive seam cell proliferation in animals homozygous for *lin-28* gain-of-function alleles that lack the 3’UTR microRNA complementary sites (*ma582* is a swap of the *lin-28* 3’UTR with a 64-nucleotide fragment of *act-1* 3’UTR, and *ma562* deletes only the *lin-4* and *let-7* complementary sites). **E:** Loss of *lin-66* results in excessive seam cell proliferation in animals homozygous for a *lin-14gf* allele, *lin-14(n355n679ts).* (*n355* is a deletion of most of the *lin-14* 3’ UTR (Wightman et al., 1993), and *n679ts* is a reduction-of-function point mutation (Reinhart & Ruvkun, 2001; Z. Shi et al., 2013)). **F:** Loss of *lin-66* results in excessive seam cell proliferation in *lin-28(ma562::GFP);lin-14(n355n679ts)* mutants that lack the microRNA-response elements of both *lin-28* and *lin-14*. **G:** Knockdown of *lin-66* exacerbates extra seam cell proliferation phenotypes of *lin-28(lf)* and *lin-46(lf)* single and compound mutants developing through L2d (induced by the temperature-sensitive *daf-7(e1372)* allele). The Student’s t test is used to calculate statistical significance (P): n.s. (not significant) p > 0.05, *p < 0.05, **p < 0.01, ***p < 0.001, ****p < 0.0001

To assess how key heterochronic regulators contribute to the *lin-66(lf)* seam cell phenotype, we knocked down *lin-14* and *hbl-1* in the *lin-66(lf)* background (Fig. 3B) and generated compound mutants with *lin-28* and *lin-46* (Fig. 3C). Knockdown of *lin-14* in *lin-66(lf)* mutants produced a phenotype resembling *lin-14* loss of function alone, indicating that *lin-14* activity is required for the seam cell overproliferation caused by loss of *lin-66*. By contrast, despite the role of *hbl-1* in promoting proliferative seam cell divisions (Abrahante, Daul, Li, Volk, Tennessen, Miller, & Rougvie, 2003; Lin et al., 2003), reducing *hbl-1* by RNAi did not suppress the seam cell overproliferation caused by loss of *lin-66*. Consistent with previous work, *lin-28(lf)* also suppressed *lin-66(lf)* seam cell phenotype, suggesting that misexpression of LIN-28 contributes to *lin-66(lf)* defects. However, because LIN-14 and LIN-28 participate in a reciprocal positive regulatory module (Fig. 3A) (Moss et al., 1997; Arasu et al., 1991; Tsialikas et al., 2017), these epistasis results alone cannot distinguish whether *lin-66* regulates LIN-14 and LIN-28 independently or whether one of these factors is regulated directly by LIN-66, and altered expression of that relatively direct target secondarily affects the other.

To test whether *lin-66* acts solely through the previously described *lin-28/lin-46* axis, we compared seam cell phenotypes of *lin-28(lf);lin-66(lf);lin-46(lf)* triple mutants to *lin-28(lf);lin-46(lf). lin-28(lf);lin-66(lf);lin-46(lf)* triple mutants displayed a significant increase in seam cell numbers compared to *lin-28(lf);lin-46(lf)* (Fig. 3C), indicating that the loss of *lin-66* can promote seam cell overproliferation even when *lin-28/lin-46* axis is absent. Moreover, *lin-66(lf);lin-46(lf)* double mutants had substantially more seam cells than either single mutant. Together, these results indicate that although *lin-28* and *lin-14* contribute to the *lin-66(lf)* phenotype, *lin-66* does not act solely through the *lin-28/lin-46* axis and may instead function in parallel.

### *lin-66* functions independently of the microRNA-binding sites in the *lin-28* and *lin-14* 3’UTRs

Previous transgene-based experiments suggested that *lin-66* repression on *lin-28* requires the *lin-28* 3’UTR (Morita and Han, 2006). We therefore asked whether the native *lin-28* 3’UTR is required for *lin-66* function in seam cell fate patterning. To address this, we examined animals carrying *lin-28(ma582)*, an allele in which the native *lin-28* 3’ UTR is replaced with a 64-nucleotide fragment of *act-1* 3’UTR (Nelson & Ambros, 2025). If *lin-66* acts primarily through the native *lin-28* 3’ UTR, then *lin-28(ma582);lin-66(lf)* doubly-mutant animals would be expected to phenocopy *lin-28(ma582)* or *lin-66(ok3326)* single mutants. However, *lin-28(ma582);lin-66(lf)* double mutants displayed pronounced seam cell hyperproliferation, with seam cell numbers significantly exceeding those of either single mutant (Fig. 3D). Similarly, *lin-28(ma562);lin-66(lf)* animals, in which the *lin-4* and *let-7-family* microRNA complementary sites are deleted from the *lin-28* 3’ UTR (Nelson & Ambros, 2025), showed enhanced seam cell hyperproliferation beyond either single mutant (Fig. 3D), indicating that *lin-66* function in seam cell fate regulation is independent from the *lin-28* 3’ UTR microRNA-responsive elements. Strikingly, the hyperproliferating seam cells in *lin-28(ma562);lin-66(lf)* mutants showed abundant LIN-14::GFP expression even at the late adult stage, when LIN-14::GFP is not detected in wild-type worms (Supp. Fig. S1). Consistent with this failure to extinguish temporal regulator expression at later larval stages, *lin-28(ma562);lin-66(lf)* adults fail to express the adult hypodermal collagen marker *Pcol-19::GFP* (Supp. Fig. S2).

To examine the relative contributions of *lin-28*, *lin-14*, and *hbl-1* to the seam cell hyperproliferation observed in *lin-28(ma562);lin-66(ok3326)* mutants, we individually knocked down each gene in the *lin-28(ma562);lin-66(ok3326)* background. Knockdown of each gene significantly suppressed the *lin-28(ma562);lin-66(ok3326)* hyperproliferation phenotype (Fig. 5A), indicating that all three regulators contribute to excessive seam cell proliferation in *lin-28(ma562);lin-66(ok3326)* animals. Knockdown of *hbl-1* and *lin-14* produced stronger suppression than *lin-28*, suggesting that misexpression of *hbl-1* and *lin-14* may be major drivers of seam cell hyperproliferation in this background. However, differences in RNAi efficiency or in the timing of knockdown relative to developmental progression could also contribute to the different degrees of suppression.

Because knockdown of *lin-14* suppressed *lin-66(lf)* (Fig. 3B), and LIN-14::GFP expression persisted in *lin-28(ma562);lin-66(ok3326)* mutants in later-larval/adult stages (Supp. Fig. S1), we examined whether *lin-66* function depends on cis-regulatory elements in the *lin-14* 3’UTR. We used the *lin-14(n355n679)* allele, which combines *n355,* a deletion removing most of the *lin-14* 3’UTR, including the *lin-4* microRNA complementary sites, with the temperature-sensitive loss-of-function mutation *n679ts* (Reinhart & Ruvkun, 2001; Shi et al., 2013; Wightman et al., 1991). Loss of *lin-66* in the *lin-14(n355n679)* background increased seam cell proliferation (Fig. 3E), indicating that *lin-66* function in seam cell patterning does not depend exclusively on the *lin-14* 3’ UTR.

The above findings, where *lin-66(lf)* could cause extra seam cell proliferation defects in the absence of the *lin-28* 3’ UTR (Fig. 3D) or in the absence of most of the *lin-14* 3’ UTR (Fig. 3E) suggests that *lin-66* could regulate *lin-14* and *lin-28* by mechanisms independent of the microRNA regulation of these genes (for example by binding to the *lin-14* or *lin-28* transcripts outside of their 3’ UTRs). However, these experiments did not exclude the possibility that *lin-66* could negatively regulate both *lin-14* and *lin-28* via their 3’ UTRs such that de-repression of either LIN-14 or LIN-28 is sufficient to produce extra seam cell numbers in *lin-66(lf).* To test this possibility, we examined the effect of *lin-66(lf)* in animals carrying both *lin-28(ma562)* and *lin-14(n355n679),* which results in simultaneous disruption of microRNA-mediated regulation of both *lin-14* and *lin-28*, and causes strong seam cell overproliferation. Importantly, loss of *lin-66* further enhanced the extra seam cell phenotype of the *lin-28(ma562);lin-14(n355n679ts)* (Fig. 3F), indicating that *lin-66* can limit seam cell proliferation independently of the microRNA binding sites in the *lin-28* and *lin-14* 3’UTRs.

### *lin-66* is required for proper seam cell fate in both continuous and L2d/dauer interrupted developmental trajectories

The heterochronic network is modulated by the worm’s developmental trajectory—whether it proceeds continuously or through the L2d/dauer-interrupted trajectory (Euling & Ambros, 1996; Ilbay & Ambros, 2019a; Karp & Ambros, 2012; Z. Liu & Ambros, 1991). We therefore asked whether the requirement for *lin-66* in seam cell fate patterning depends on developmental trajectory. To address this, we used the temperature-sensitive *daf-7(e1372)* mutant, which constitutively forms dauers at 25 °C but develops through the L2d stage at 20 °C. At 20 °C, *lin-66* knockdown in the *daf-7(e1372)* background increased seam cell numbers to a degree similar to that observed in wild-type animals (Fig. 3G), indicating that *lin-66* is required for proper seam cell patterning during both continuous and L2d-interrupted development.

Consistent with previous work, loss of *lin-46* in the *daf-7(e1372)* background resulted in extra seam cells, supporting a critical role for *lin-46* during L2d-interrupted development (Karp and Ambros 2012). *lin-66* knockdown further enhanced *lin-46(lf)* seam cell phenotype in both L2d-interrupted and continuous developmental trajectories (Fig. 3G). The effect of *lin-66* knockdown appeared more pronounced in the *daf-7(e1372)* backgrounds than in the corresponding continuously developing strains, suggesting that *lin-66* may become particularly important when seam cell fate progression occurs through the L2d associated trajectory.

### Loss of function of GW182 homologs *ain-1* and *ain-2* enhances the seam cell fate defects of lin-66(lf)

LIN-66 was previously recovered by co-immunoprecipitation with the *C. elegans* GW182 family protein AIN-1, suggesting that LIN-66 may function in conjunction with miRISC-mediated regulation (Mayya et al., 2021). We therefore examined the genetic relationship between *lin-66* and the partially redundant GW182-homologs *ain-1* and *ain-2.* The *ain-1(ku322)* null mutants display slightly elevated seam cell numbers and incomplete adult alae formation– consistent with a slight disruption in heterochronic gene cascade microRNA activity– while *ain-2(tm1863)* reduction-of-function mutants exhibit essentially normal seam cell counts and no alae defects (Ding et al., 2005; Zhang et al., 2007). By contrast, *ain-2(tm1863)*;*ain-1(ku322)* double mutants exhibit severe heterochronic defects with dramatically increased seam cell numbers (more than 50 seam nuclei and no alae reported in adults), reflecting the partially redundant functions of the two paralogs in miRISC activity (Zhang et al., 2007).

Strikingly, *lin-66(ok3326)*; *ain-1(ku322)* double mutants displayed substantially greater seam cell hyperproliferation than either single mutant (Fig. 4A). This enhancement indicates that *lin-66* and *ain-1* function in parallel to restrain reiterative seam cell divisions. To assess whether the second GW182 paralog functions in a similar capacity, we combined *lin-66(ok3326)* with the *ain-2(tm2432)* loss-of-function allele (Fig. 4B). *ain-2(tm2432);lin-66(ok3326)* resulted in a stronger extra-seam cell proliferation phenotype than either mutation alone. Thus, loss of either GW182 homolog sensitizes seam cell-fate patterning to *lin-66(lf)*, consistent with LIN-66 acting in parallel with GW182-dependent gene regulation.

**Figure 4:**
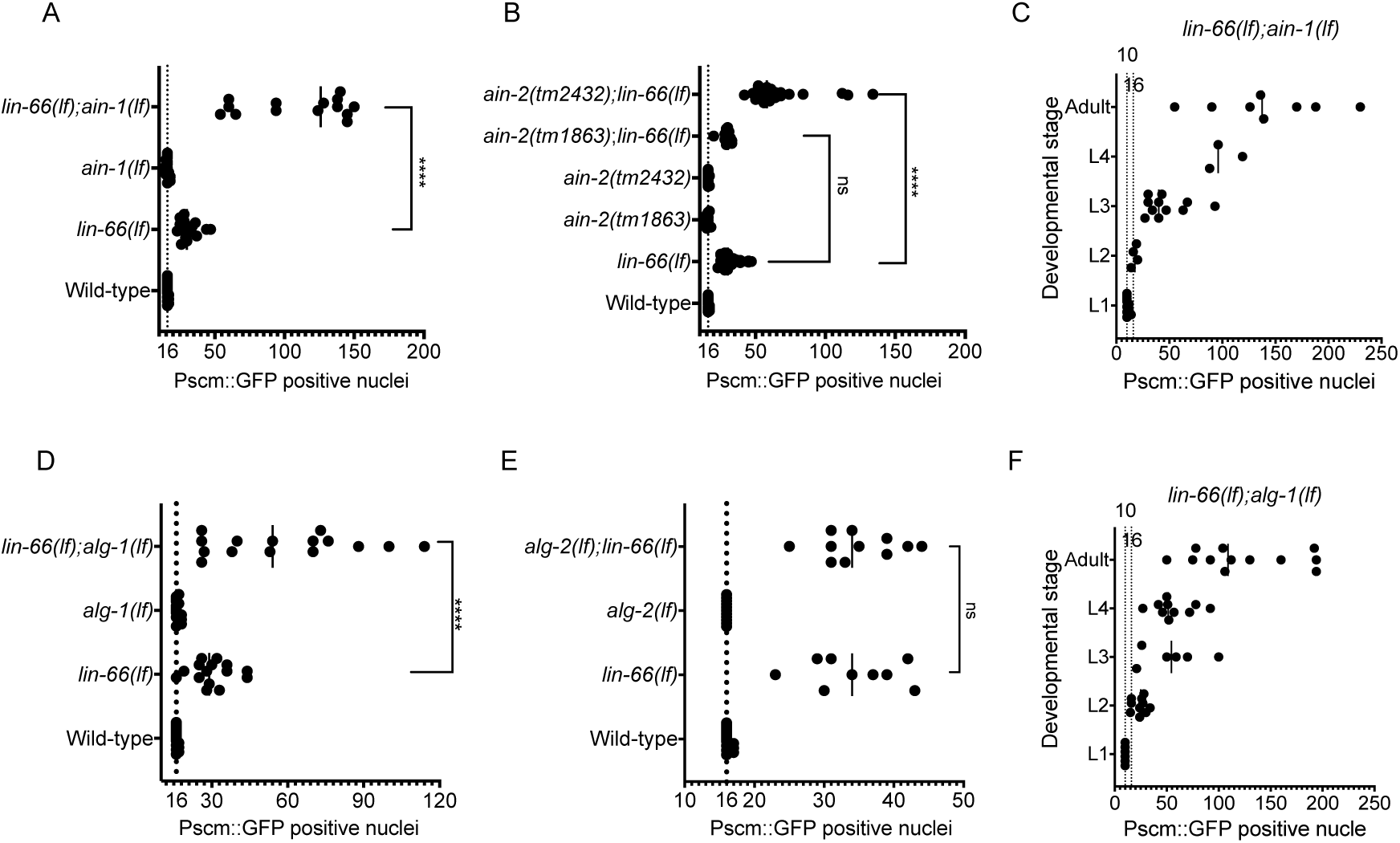
loss of GW182-like genes *ain-1* and *ain-2,* and the miRISC-Argonaute gene *alg-1* enhances the extra seam cell defect of *lin-66(lf).* **A:** *ain-1(lf)* significantly enhances the *lin-66(lf)* extra seam cell defects, resulting in hyperproliferation phenotype. B: *ain-2(tm2432)* (a presumed null) aggravates the seam cell phenotype of *lin-66(lf)* but *ain-2(tm1863)* (a presumed reduction of function allele) does not. *alg-1(lf)* D, but not *alg-2(lf)* E aggravates the seam cell phenotype of *lin-66(lf)*. **C, F:** Number of Pseam::GFP positive nuclei in *lin-66(lf);ain-1(lf)* and a*lg-1(lf);lin-66(lf)* mutants progressively increase throughout larval development. The Student’s t test is used to calculate statistical significance (P): n.s. (not significant) p > 0.05, *p < 0.05, **p < 0.01, ***p < 0.001, ****p < 0.0001

Interestingly, when *lin-66(lf)* was combined with a reduction-of-function (*rf*) allele of *ain-2*, *tm1863*, the *ain-2(tm1863);lin-66(ok3326)* double mutants were indistinguishable from *lin-66(ok3326)* single mutant animals (Fig. 4B). This suggests that *ain-2(tm1863)* retains a level of residual AIN-2 activity that is sufficient to limit seam cell overproliferation in the absence of *lin-66*.

The genetic interaction between *lin-66* and the AIN paralogs also extend beyond seam cell patterning. Although *ain-2(rf);ain-1(lf)* mutants are viable, their viability depends on residual AIN-2 activity, as further depletion of AIN-2 by RNAi renders them nonviable (Zhang et al., 2007). Similarly, we were unable to recover viable *ain-2(tm1863);lin-66(ok3326);ain-1(lf)* animals. Thus, loss of *lin-66* further compromises a sensitized background in which GW182 activity is limiting, suggesting that LIN-66 may support developmental robustness when miRISC effector function is reduced.

### Loss of miRISC Argonaute gene *alg-1,* but not *alg-2* enhances the seam cell fate defects of lin-66(lf)

Because *lin-66* showed strong genetic interactions with the GW182-like genes *ain-1* and *ain-2*, we examined the functional relationship of *lin-66* to the microRNA Argonaute genes *alg-1* and *alg-2*. Consistent with previous observations (that employed *lin-66(ku423)* and *alg-1(gk214)* loss-of-function alleles) (Morita & Han, 2006), *lin-66(ok3326);alg-1(tm492)* double mutants displayed dramatically increased seam cell numbers compared with either single mutant (Fig. 4D). This enhancement indicates that *lin-66* and *alg-1* may function in parallel to limit reiterative seam cell divisions. By contrast, *alg-2(ok304);lin-66(ok3326)* double mutants resembled *lin-66(ok3326)* single mutants (Fig. 4E), suggesting that *lin-66* and *alg-2* may not share downstream targets in the heterochronic gene cascade.

We next asked which heterochronic regulators drive the hyperproliferation observed when both *lin-66* and *ain-1* functions are compromised. In *lin-66(ok3326);ain-1(ku322)* mutants, *lin-14* knockdown strongly suppressed seam cell hyperproliferation, whereas *lin-28* and *hbl-1* knockdowns had little effect (Fig. 5B). This pattern contrasts with the suppression profile of *lin-28(ma562);lin-66(ok3326)* mutants (Fig. 5A) (which are most effectively suppressed by *hbl-1* knockdown) and suggests that the *lin-66(ok3326);ain-1(ku322)* hyperproliferation phenotype is driven primarily by persistent LIN-14 activity.

**Figure 5:**
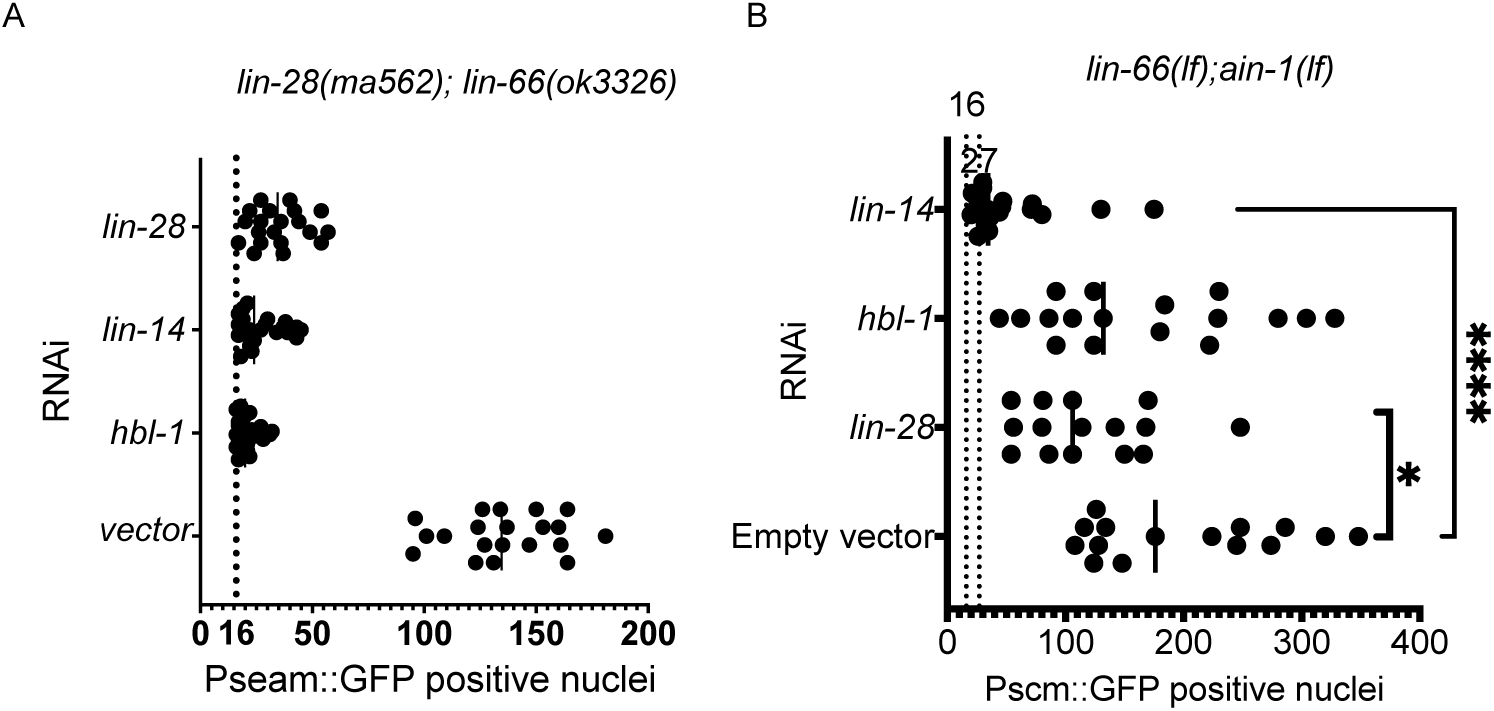
Seam cell hyperproliferation phenotype of *lin-28(ma562);lin-66(lf)* and *lin-66(lf);ain-1(lf)* and mutants upon knockdown of *lin-14*, *lin-28,* and *hbl-1*. A: Excessive seam cell proliferation in *lin-28(ma562gf);lin-66(lf)*compound mutants is partially suppressed by knockdown of *lin-28*, *lin-14*, or *hbl-1*. Knockdown of *hbl-1* results in the most potent suppression, followed by *lin-14* and *lin-28.* B: Hyperproliferation phenotype of *lin-66(lf);ain-1(lf)* is suppressed weakly by *lin-28* knockdown and strongly by *lin-14* knockdown.

### *lin-66(lf);alg-1(lf)* double mutants exhibit synthetic early developmental arrest and lethality

The interaction between *lin-66* and *alg-1* extended beyond seam cell patterning. Whereas *lin-66(ok3326)* and *alg-1(tm492)* single mutants each showed approximately 30% embryonic or early larval arrest, *lin-66(ok3326);alg-1(tm492)* double mutants exhibited pronounced synthetic lethality, with approximately 70% of animals failing to hatch or develop past the L1 stage (Fig. 6A). Thus, *lin-66* becomes especially vital when *alg-1-*dependent microRNA activity is reduced, supporting a broader role for *lin-66* in reinforcing miRISC-regulated developmental programs.

**Figure 6:**
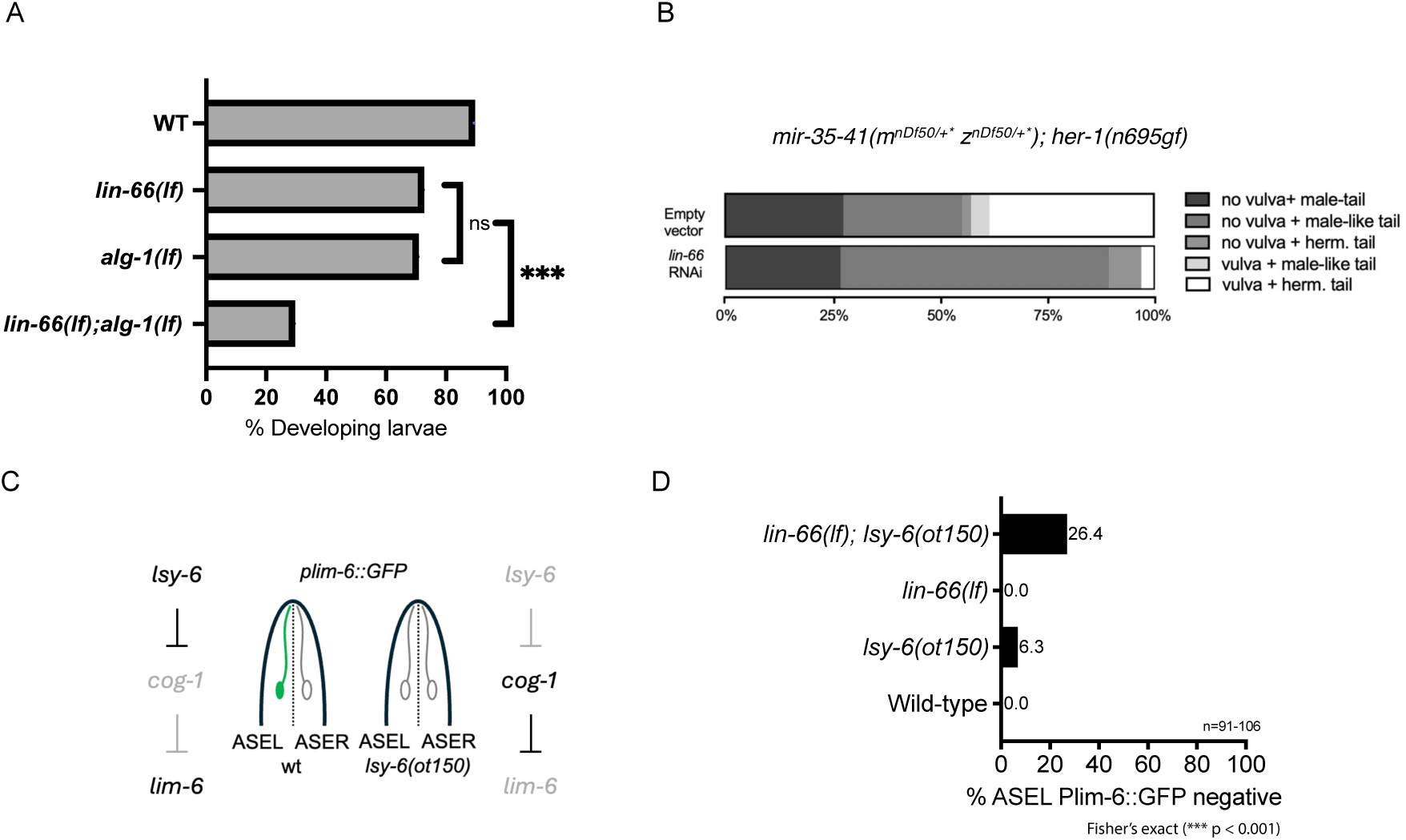
**A:** *lin-66(lf)* enhances the embryonic and larval arrest phenotype of *alg-1(lf).* Approximately 70 % of embryos from *lin-66(lf)* hermaphrodites or *alg-1(gk214)* hermaphrodites develop through larval stages by 7 days after plating, while only 30% of embryos from doubly-mutant *lin-66(ok3326);alg-1(gk214)* hermaphrodites complete larval development by 7 days (n=700-900). **B:** *lin-66* modifies the dose-dependent sex determination process that is governed by *mir-35-41*. Knock down of *lin-66* results in masculinization of the *mir-35-41(mnDf50/+* znDf50/+*);her-1(n695gf)*. Loss of *lin-66* enhances the ASER/ASEL cell fate defect of *lsy-6(ot150)*. **C:** Diagram of *C. elegans* head illustrating the ASEL and ASER neurons. *Isy-6* microRNA expression in ASEL mediates the asymmetric expression of Plim-6::GFP reporter in ASEL. Animals homozygous for *lsy-6(ot150)* can exhibit absence of Plim-6::GFP reporter in the ASEL neuron. **D:** Quantification of Plim-6::GFP expression phenotypes of *Isy-6* and *lin-66* single and compound mutants.

### *lin-66* influences the ASER/ASEL cell fate specification determined by microRNA *lsy-6*

To ask whether *lin-66* modulates microRNA-regulated developmental decisions outside the heterochronic pathway, we used a sensitized assay for *lsy-6* microRNA function in ASE neuronal fate specification. The *lsy-6* microRNA is essential during embryogenesis for specifying the distinct identities of the bilaterally symmetric ASE sensory neurons, ASEL and ASER (Fig. 6C). Specifically, *lsy-6* mediates this asymmetry by repressing its target, *cog-1*, exclusively in ASEL neurons, thereby initiating a cascade of gene expression events critical for proper neuronal fate determination (Johnston & Hobert, 2003). To sensitively assess the functional involvement of *lin-66* in the ASEL/R fate, we utilized the *lsy-6(ot150)* hypomorphic allele, which reduces, but does not eliminate, *lsy-6* expression, creating a scenario sensitive to perturbations in microRNA activity (Sarin et al., 2007). In this assay, *lim-6p::GFP* expression marks ASEL fate and provides a readout of successful *lsy-6-*dependent repression of *cog-1*.

Under our conditions, *lsy-6(ot150)* animals showed a low penetrance of ASEL fate defect, with ∼6.3% of the population lacking wild-type *lim-6p::GFP* expression pattern (Fig. 6D). *lin-66(lf)* single mutants showed no detectable ASEL cell fate defects, indicating that *lin-66* is not essential for ASEL specification when *lsy-*6 function is intact. However, *lin-66(lf);lsy-6(ot150)* double mutants showed a marked increase in ASE cell fate defect, with 26.4% of animals affected. Thus, loss of *lin-66* sensitizes animals to reduced *lsy-6* activity, suggesting that *lin-66* contributes to the robustness of microRNA-regulated cell fate decisions.

### *lin-66* knockdown exacerbates the cryptic masculinization of *mir-35-41(nDf50)*

To further assess the involvement of *lin-66* in microRNA-regulated pathways, we investigated its role in sex determination, a process regulated by *mir-35* family microRNAs (McJunkin & Ambros, 2017). We examined interactions between *lin-66* and the *mir-35* family in the context of *her-1(n695gf)*, a gain-of-function mutation in the gene encoding HER-1, the master driver of male sexual fate. *her-1(n695gf)* causes mild masculinization in XX animals, resulting in pseudomales with male phenotypic features despite having XX chromosomes. Previous work has established that loss of the *mir-35* family microRNAs dramatically enhances this masculinization phenotype in a dosage-dependent manner (McJunkin & Ambros, 2017). Since the *mir-35-41(nDf50);her-1(n695gf)* double mutant produces nearly completely masculinized, self-sterile XX pseudomales with high penetrance, we utilized the more experimentally tractable *mir-35-41(nDf50/+);her-1(n695gf)* heterozygotes. When these heterozygous hermaphrodites were subjected to *lin-66* RNAi from the L4 stage, their F1 progeny -- *mir-35-41(nDf50/+);her-1(n695gf)*; *lin-66(RNAi)* -- exhibited dramatically enhanced masculinization compared to animals treated in parallel with control RNAi (Fig. 6B). This synthetic enhancement of the masculinization phenotype suggests that *lin-66* facilitates the *mir-35* family function in repressing targets involved in sex determination, providing additional evidence for *lin-66*’s broader role in conjunction with microRNA-mediated regulation across different physiological contexts.

## DISCUSSION

### Functional interactions of *lin-66* with microRNA regulatory processes

Previous studies of *C. elegans lin-66(lf)* mutants reported heterochronic developmental phenotypes and ectopic LIN-28 protein expression, suggesting that *lin-66* function intersects with post-transcriptional regulation in the heterochronic pathway (Morita & Han, 2006). The recovery of LIN-66 in association with AIN-1, NHL-2 and NTL-1 (Mayya et al., 2021; Wu et al., 2016) further suggested that LIN-66 may act in or near miRISC-associated regulatory complexes. Here, using genetic epistasis and fluorescent reporter assays, we show that *lin-66* limits inappropriate developmental persistence of the early temporal regulators LIN-14 and LIN-28.

Consistent with functional overlap between LIN-66 and microRNA-mediated regulation, *lin-66(lf)* enhanced the heterochronic phenotypes of animals genetically-sensitized for miRISC activity, by mutations of *alg-1* or the GW182-like genes *ain-1* and *ain-2* (Figs. 4D, 4A, 4B). The genetic interactions of *lin-66* with Argonaute genes was not the same for both paralogs: *alg-1(lf)*, but not *alg-2(lf)*, strongly enhanced *lin-66(lf)* seam-cell defects (Figs. 4D&4E), suggesting that the relationship between LIN-66 and microRNA activity may be context or complex-specific. *lin-66(lf);alg-1(lf)* also exhibited synthetic embryonic and early larval arrest (Fig. 6A), indicating that LIN-66 becomes particularly important when ALG-1-dependent microRNA activity is compromised. This genetic relationship is consistent with broader evidence that CSD proteins can functionally engage Argonaute-associated pathways; for example, CSDE1 was reported to interact with AGO2-containing miRISC and contribute to miRNA-mediated silencing (Kakumani et al., 2020).

The genetic interaction between *lin-66* and microRNA-mediated regulation extended beyond the heterochronic pathway. In sensitized non-hypodermal contexts, *lin-66(lf)* enhanced the ASE neuronal fate defects of *lsy-6(ot150)* animals (Fig. 6C and 6D), and *lin-66* RNAi enhanced the masculinization phenotype of *mir-35-41/+;her-1(gf)* animals (Fig. 6B). Together, these results suggest that LIN-66 supports the robustness of multiple microRNA-sensitive developmental programs, rather that acting solely within the heterochronic pathway.

### *lin-66(lf)* dysregulates the *lin-14/lin-28* nexus of the heterochronic cascade

Previous epistasis placed *lin-66* upstream of *lin-28* (Morita & Han, 2006). Consistent with this relationship, we found that *lin-28(lf);lin-66(lf)* animals resemble *lin-28(lf)* single mutants in seam cell phenotype (Fig. 3C). Previous western blot analysis showed that LIN-28 protein persists at later larval stages in *lin-66(lf),* and our analysis of endogenously tagged LIN-28:GFP similarly revealed modest persistence after the L2 stage (Fig. 2A). Thus, *lin-66* contributes to the developmental downregulation of LIN-28.

However, *lin-66* also affected seam cell proliferation independently of the *lin-28;lin-46* axis: loss of *lin-66* increased seam cell number even in *lin-28(lf);lin-46(lf)* background (Fig. 3B). This finding indicates that LIN-28 misregulation is not the only contributor to the *lin-66(lf)* seam cell phenotype. Our data point to LIN-14 as an additional, and likely major, contributor. Knockdown of *lin-14* suppressed *lin-66(lf)* seam cell overproliferation (Fig. 3B), and LIN-14::GFP persisted ectopically after the L1 stage in *lin-66(lf)* animals (Fig. 2B). Moreover, LIN-14::GFP remained abundant in late larval and adult-stage *lin-28(ma562);lin-66(ok3326)* animals, whereas it was much more strongly downregulated in *lin-28(ma562)* single mutants (Supp. Fig. S1). By contrast, *hbl-1* RNAi did not suppress the *lin-66(lf)* seam cell phenotype, and HBL-1 downregulation was not detectably perturbed in *lin-66(lf)* mutants (Fig. 2C). Together, these results support a model in which *lin-66* limits inappropriate persistence of early temporal regulators, with LIN-14 misregulation making a major contribution to the heterochronic phenotype caused by loss of *lin-66.* Prior genetic analysis of *lin-14* supports a dose- and stage-dependent model for temporal fate specification, in which high LIN-14 activity promotes L1-specific programs, reduced LIN-14 activity is compatible with L2-stage fates, and further downregulation after L2 is required for later larval programs (Ambros & Horvitz, 1987). In addition, structure-function analysis of LIN-14 supports separable activities associated with distinct LIN-14 isoforms that differ in their amino-terminal sequences (Reinhart & Ruvkun, 2001), raising the possibility that persistent LIN-14 in *lin-66(lf)* animals could affect stage-specific seam cell programs in an isoform-dependent manner. However, we do not yet know which LIN-14 isoform is misregulated in *lin-66(lf)* mutants, or whether LIN-66-dependent regulation is isoform-specific. It also remains unclear whether the seam cell hyperproliferation phenotype seen in various *lin-66(lf)* compound mutants reflects repetition of L2 fates, or a more complex failure to achieve larva-to-adult seam lineage fates.

### *lin-66*-dependent regulation is separable from canonical 3’UTR-mediated microRNA repression

Although LIN-66 has been reported to co-immunoprecipitate with AIN-1, NHL-2, and NTL-1, it remains unclear whether these interactions are direct or RNA-mediated. We therefore asked whether LIN-66-dependent regulation requires 3’UTR microRNA-responsive elements that mediate canonical miRISC regulation of *lin-14* and *lin-28.* Loss of *lin-66* enhanced seam cell proliferation when either the native *lin-28* or *lin-14* 3’UTRs were largely deleted (Figs. 3D-F). Moreover, loss of *lin-66* enhanced the extra seam cell phenotype of *lin-28(ma562);lin-14(n355n679ts)* animals, in which canonical 3’UTR-mediated microRNA regulation of both genes is disrupted. Thus, the canonical microRNA-binding sites in the *lin-28* and *lin-14* 3’UTRs are not required for expression of *lin-66(lf)* extra seam cell phenotype.

These findings argue that LIN-66-dependent regulation is separable from canonical 3’UTR-mediated microRNA repression. If LIN-66 acts directly on *lin-28* or *lin-14* transcripts, the relevant regulatory information likely lies outside the canonical microRNA-binding sites. Alternatively, LIN-66 may indirectly affect LIN-14 and LIN-28 expression. Although we cannot exclude the effects on microRNA biogenesis in all contexts, previous work found that levels of the major heterochronic microRNAs *lin-4, mir-48* and *mir-84* were unchanged in *lin-66(lf)* mutants (Morita & Han, 2006), arguing against a broad defect in microRNA biogenesis.

### Conserved residues in the LIN-66 CSD are required for heterochronic function

Cold shock domain-containing proteins bind RNA and DNA through conserved stacking interactions between nucleic acid bases and aromatic residues, a mechanism that is often with limited sequence specificity (Heinemann & Roske, 2021). Previous studies have identified four conserved ssRNA/ssDNA binding motifs among CSD proteins (RNP1 in β2 strand, RNP2 in β3 strand, KWFN in β1 strand, and EGFKTL/ EGFRSL in the β3β4 loop) (Kleene, 2018). Three of the four conserved CSD residues that we mutated lie within these predicted motifs (Fig. 1C). LIN-66(CSD*) showed a subcellular expression pattern similar to wild-type LIN-66 but was not functional in the context of seam cell patterning (Fig. 1F). This suggests that the potential RNA-binding activity of LIN-66 is not essential for its localization to subcellular puncta, but that puncta localization alone is insufficient for its function in seam cell fate patterning.

The molecular basis of LIN-66 activity remains to be determined. Future studies will be required to identify *in vivo* LIN-66 target RNAs and define whether LIN-66 affects transcript abundance, localization, or translational efficiency. To date, multiple CSD-containing proteins have been shown to modulate microRNA-related pathways through diverse mechanisms. In *C. elegans,* the CSD-containing protein LIN-28 binds primary *let-7* transcripts to inhibit *let-7* processing, and *in vivo* HITS-CLIP further revealed LIN-28 interactions with *let-7* precursors and broader mRNA networks (Stefani et al., 2015; Van Wynsberghe et al., 2011). Similarly, mammalian LIN28 uses its CSD together with its zinc-finger domain to bind *pre-let-7* and regulate *let-7* biogenesis (Büssing et al., 2008; Mayr et al., 2012; Nam et al., 2011; Ustianenko et al., 2018). Moreover, mammalian YBX1 can bind specific miRNAs, including miR-223, through its CSD and promote their sorting into exosomes together with YBAP1 (X.-M. Liu et al., 2021; Ma et al., 2023). CSDE1 provides another example of an AGO2-specific CSD protein that functions in microRNA pathways, promoting Dicer-independent miR-451 maturation and enhancing target decay by recruiting elements of the decapping machinery (Kakumani et al., 2020, 2023). These examples illustrate that CSD proteins can influence microRNA-related pathways at multiple levels. The combination of a CSD and extensive intrinsically disordered regions raises the possibility that LIN-66 participates in context-dependent RNA-protein assemblies, which may help explain why LIN-66 has been associated with translational activation in motor neurons (Blazie et al., 2024a) but with reduced accumulation of LIN-14 and HBL-1 in hypodermal cells.

## MATERIALS AND METHODS

### *C. elegans* culture conditions

The *C. elegans* strains used in this study are listed in Table S1. Strains were maintained at 20 ^0^C on nematode growth media (NGM) seeded with *E.coli* HB101 strain unless otherwise indicated.

### Egg preparation and L1 synchronization

Egg-laying/or gravid adults from NGM plates were washed and collected in 15 ml conical tubes with M9 media. Worms were pelleted with a clinical centrifuge. The worm pellets were treated with alkaline bleach solution (0.5 M NaO, 1% sodium hypochlorite) for 6 minutes with constant shaking. Extracted embryos were pelleted and rinsed three times with M9 buffer. For L1 synchronization, extracted embryos were incubated and hatched in M9 buffer at 20 ^°^C with mild shaking in 15 mL conical tubes for ∼16-20 hours.

### RNAi knockdown

To knock down genes of interest, RNAi was performed via the feeding method (Conte et al., 2015). The RNAi vectors were designed by selecting coding sequence fragments that target all isoforms of the specific gene, and the fragments were then cloned into the T444T super RNAi vector (Sturm et al., 2018) by GenScript cloning services. RNAi vectors were transformed into the *E. coli* HT115 strain. Experiments in Fig. 3G were performed using the RNAi clones from the Ahringer library (Kamath et al., 2003). Other RNAi clones used in this study are listed in Table S2. Overnight cultures of each bacterial clone expressing dsRNA were diluted in LB broth with 100 μg/ml ampicillin and cultured at 37 ^0^C until the OD_600_ values reached between 0.6-0.8. The bacterial cultures were seeded onto NGM plates containing 100 μg/ml ampicillin and 1 mM IPTG and induced at room temperature for 24-48 hours. Synchronized L1 larvae were placed on the RNAi plates at 20 ^0^C, and were scored when they reached adulthood.

### *mir-35-42* cryptic masculinization assay

For the cryptic masculinization assay (McJunkin & Ambros, 2017), single *mir-35-41(nDf50)/mIn1 [myo-2::GFP] II; her-1(n695gf) V* L4 larvae were placed on RNAi plates. The F1s that are heterozygous for *mir-35-41* deletion (GFP+, non-dumpy animals-*mir-35-41(nDf50)/mIn1 [myo-2::GFP] II; her-1(n695gf) V*) were scored for the sex determination phenotypes.

### *lsy-6(ot150)* ASEL cell fate assay

Specification of ASEL neuronal cell fate was scored using the expression pattern of *Plim-6::GFP* reporter at L4/adult stages.

### Embryonic/larval arrest and lethality assay

Embryos were extracted with alkaline bleach solution as described above. Extracted embryos were placed on NGM plates seeded with *E. coli* HB101 at 20^°^C (∼100 embryos per plate). The number of hatched larvae was counted in the following seven days. This assay was also performed on physically extracted embryos (by puncturing gravid adults) and yielded similar results (data not shown).

### Quantification of the seam cells and statistical analysis

Adult worms were anesthetized in a drop of 0.2 mM levamisole solution on 2% agarose pads on imaging slides and covered with a coverslip. The slides were imaged with Zeiss Imager Z1 microscope that is equipped with ZEISS Axiocam 503 mono camera. The number of nuclei expressing the wIs51 (Pscm::GFP) marker on one lateral side of the animals was counted at 60x magnification. GraphPad Prism 10 was used for statistical analysis (Student’s t-test) ((p): n.s. (not significant) p > 0.05, *p < 0.05, *p < 0.01, ***p < 0.001, ****p < 0.0001) and data visualization.

### Imaging of endogenously tagged fluorescence reporters in *C. elegans* larvae

Worms were anesthetized in a drop of 0.2 mM levamisole solution on 2% agarose pads on imaging slides and covered with a coverslip. HBL-1::mScarlet-I was imaged using Zeiss Imager Z1 microscope that is equipped with ZEISS Axiocam 503 mono camera. LIN-14::GFP and LIN-28::GFP were imaged using Nikon CSU-W1 Yokogawa Spinning Disk Field Scanning Confocal microscope equipped with Prime BSI Express A22E726013 camera. Microscopic images were processed using Fiji (Schindelin et al., 2012).

### Endogenous tagging of *lin-66* with green fluorescence protein and creation of the CSD mutation

CRISPR/Cas9 ribonucleoprotein strategy was used C-terminally to tag *lin-66* with green fluorescent protein. CRISPR/Cas9 mix (Integrated DNA technologies (IDT) Alt-R™ S.p. Cas9 Nuclease (122 ng/µL), IDT Alt-R™ CRISPR-Cas9 crRNA RBcrRNA18 guide RNA (56 ng/ μl), IDT Alt-R™ CRISPR-Cas9 tracrRNA 100 ng/μl, partially single-stranded dsDNA donor template (linker::GFP with ∼35 bp homology arms-PCR amplified, purified and melted according to (Dokshin et al., 2018) (30 ng/μl), plasmid pOI124 (Ilbay & Ambros, 2019b) (40 ng/μl) containing *rol-6(su1006)* as co-injection marker) was prepared in IDT Nuclease Free Duplex Buffer. This mix was injected into the germlines of young *adult* N2 *C. elegans* hermaphrodites. Non-roller F1s from P0 plates with lots of roller animals were cloned and screened for the expected insertion using PCR (primers RB1039, 1040, and 1041(Table S3). F2 progeny with the homozygous insertion were cloned, and the genomic locus spanning the edit in these worms was genotyped using Sanger sequencing (with primers RB1071,1072). The resulting *lin-66::linker::GFP* allele was named *ma616*. The strain carrying the *ma616* allele was backcrossed twice and named VT4259.

Four conserved residues in the LIN-66 CSD were substituted with alanine in the VT4259 (*lin-66::linker::GFP*) background with a similar CRISPR/Cas9 strategy described above (IDT Alt-R™ S.p. Cas9 Nuclease (122 ng/µL), RBsgRNA28 IDT Alt-R™ CRISPR-Cas9 sgRNA (80 ng/µL), 390bp dsDNA donor template (100 ng/µL) (Twist Bioscience Gene Fragment, and dpy-10_sgRNA co-CRISPR marker (8 ng/µL)). The cloned worms were screened for the desired edit using the BtsI restriction enzyme recognition site introduced by the substitutions (amplified with RB1202 & RB1204).

## ACKNOWLEDGEMENTS

We thank members of the Ambros and Mello labs for the helpful discussion and comments on this project. Some strains were provided by the CGC, which is funded by the NIH Office of Research Infrastructure Programs (P40 OD010440).

## COMPETING INTERESTS

Authors declare no competing or financial interest.

## AUTHOR CONTRIBUTIONS

R.B. and V.A. designed and conducted the experiments, interpreted the results, and wrote the paper. V.A. administered the project and acquired the funding.

---

**Figure S1:**
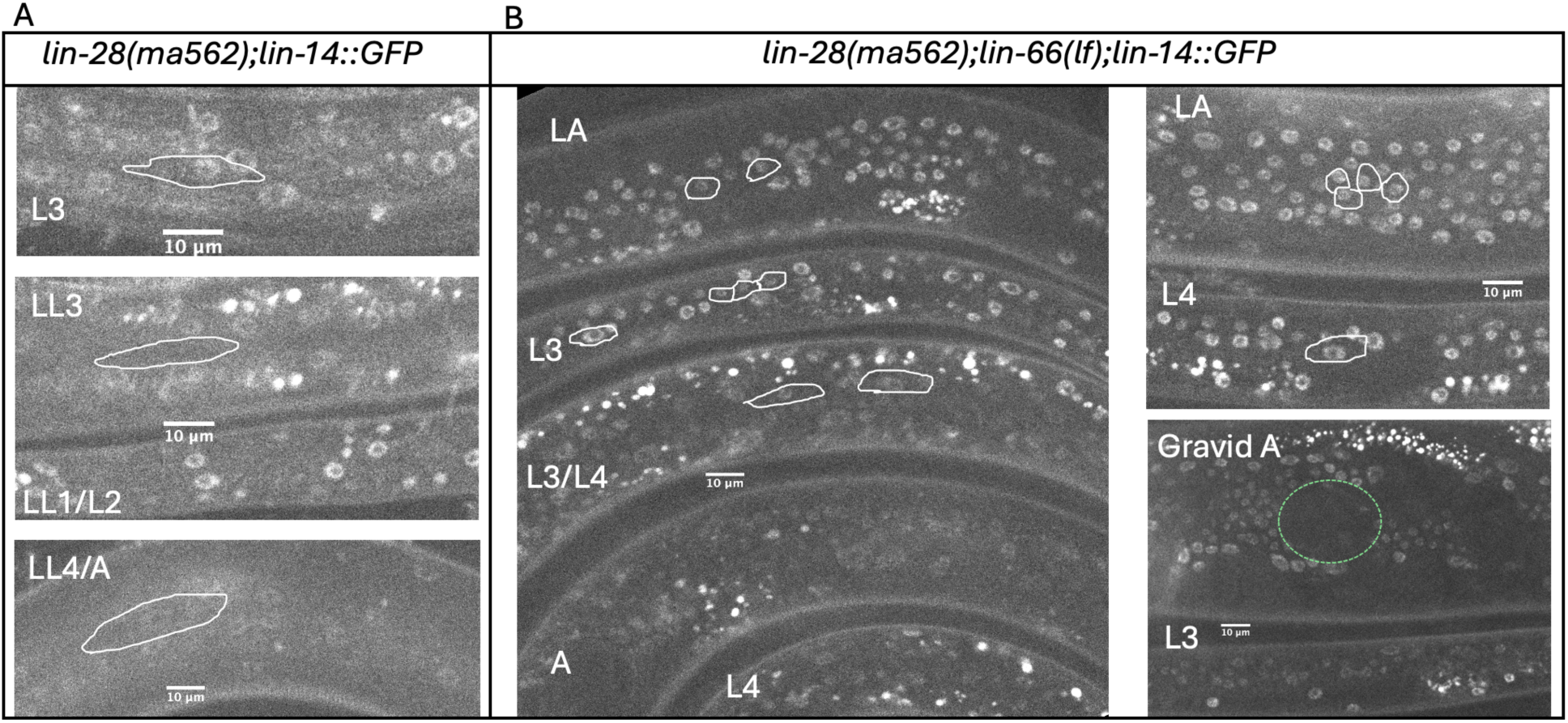
Hypodermal LIN-14::GFP expression in the wild type, *lin-28(ma562)* and *lin-28(ma562); lin-66(ok3326)* mutants. LIN-14::GFP is ectopically expressed in *lin-28(ma562); lin-66(ok3326)* mutants in later larval and adult stages. **A:** By the late L3 and L4 stages of the *lin-28(ma562)* larvae, LIN-14::GFP expression is significantly downregulated, yet still visible (example seam cells indicated by light gray borders). **B:** Abundant LIN-14::GFP expression is detected in seam cell nuclei of L3/L4 larvae and adults of *lin-28(ma562); lin-66(ok3326)*. In the Gravid adult, the shading from the embryo is circled in green dotted lines.

**Figure S2:**
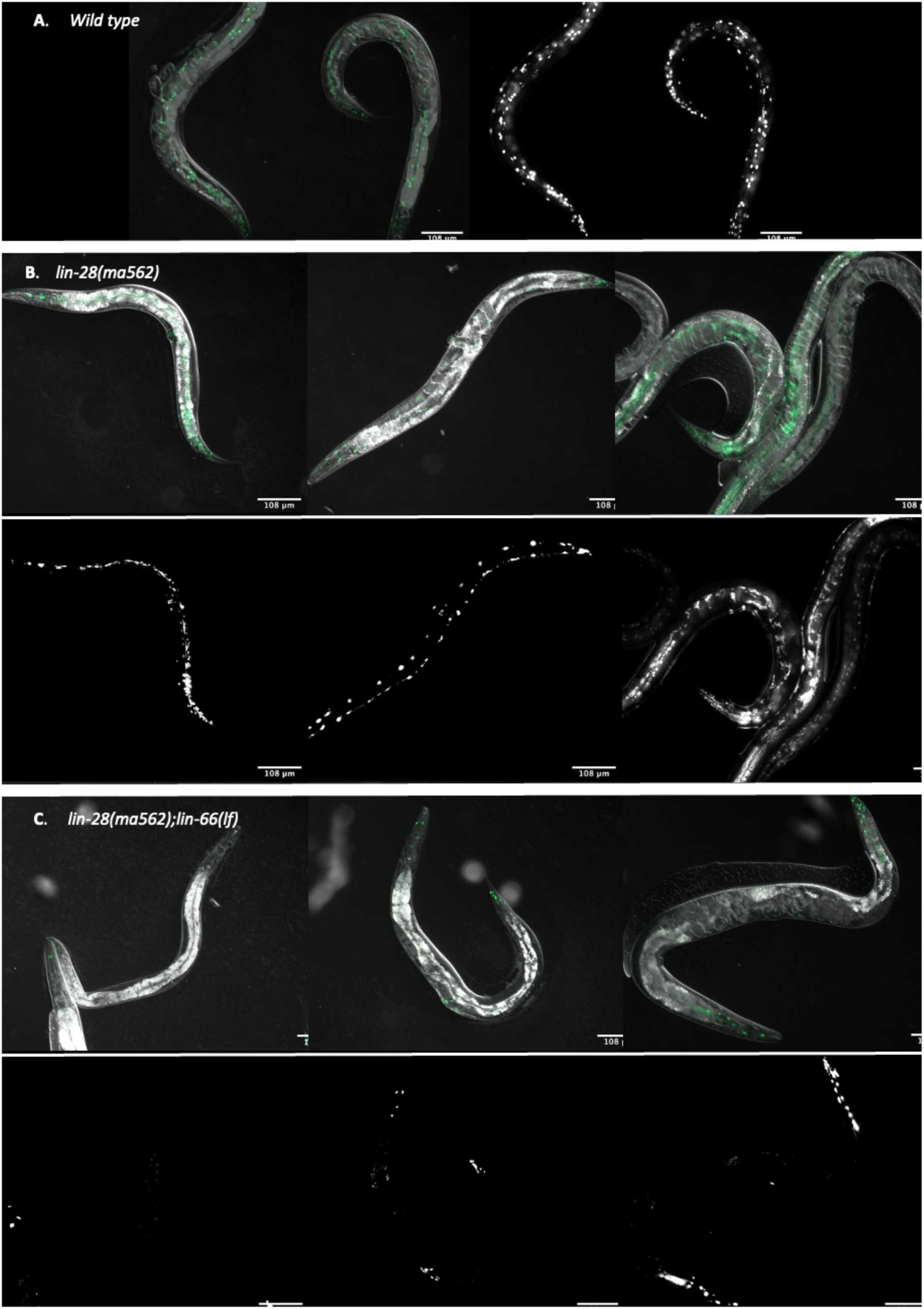
Defective adult-specific hypodermal reporter expression pattern in *lin-28(ma562);lin-66(lf)* mutants. *lin-28(ma562);lin-66(lf)* mutants are defective in expression of the adult-specific hypodermal reporter *Pcol-19::GFP*. **A:** in wild type adult hermaphrodites, *col-19::GFP* is expressed in the nuclei of hypodermal seam and hyp7 cells. Left panel, GFP overlaid on DIC image; right panel, GFP channel only. **B:** *lin-28(ma562)* adults display reduced *col-19::GFP* expression in hyp7 nuclei, with some animals expressing GFP only in seam cell nuclei (left and center) or reduced expression in seam and hyp7 (right). The upper series shows GFP overlaid on DIC; the bottom series shows the GFP channel only. **C:** *lin-28(ma562);lin-66(ok3326)* adults lack evident *col-19::GFP* in most hypodermal nuclei. Upper series shows GFP overlaid on DIC; bottom series, GFP channel only.

**Table S1:** *C. elegans* strains used in this study.

| Strain name | Genotype | Source |
| --- | --- | --- |
| VT3343 | *mir-35-41(nDf50)/mIn1 [myo-2::GFP] II; her-1(n695gf) V* | Ambros Lab |
| VT4259 | *lin-66(ma616) IV* | this study |
| VT4489 | *lin-66(ma616) IV; wIs51 V* | this study |
| VT4490 | *lin-66(ma616ma656) IV; wIs51 V* | this study |
| VT4346 | *lin-66(ok3326) IV; wIs51 V* | this study |
| VT3455 | *wIs51 V.* | Ambros Lab |
| OH3646 | *otIs114 [lim-6pro::GFP rol-6(su1006sd)] I; lsy-6(ot150)* | CGC |
| VT4491 | *otIs114 [lim-6pro::GFP rol-6(su1006sd)] I* | this study |
| VT4492 | *otIs114 [lim-6pro::GFP rol-6(su1006sd)] I; lin-66(ma646lf) IV* | this study |
| VT4493 | *otIs114 [lim-6pro::GFP rol-6(su1006sd)] I; lin-14(n179) X.* | this study |
| VT4494 | *otIs114 [lim-6pro::GFP rol-6(su1006sd)] I; lsy-6(ot150); lin-66(ma646lf) IV* | this study |
| VT4394 | *otIs114 [lim-6pro::GFP rol-6(su1006sd)] I; lsy-6(ot150); lin-14(n179) X.* | this study |
| VT4366 | *otIs114 [lim-6pro::GFP rol-6(su1006sd)] I; lsy-6(ot150); lin-66(ma646lf) IV; lin-14(n179) X.* | this study |
| VT4367 | *lin-66(ok3326) IV; wIs 51 V; alg-1(tm492) X* | this study |
| VT4495 | *wIs 51 V; alg-1(tm492) X* | this study |
| VT4357 | *alg-2(ok304) II; wIs51 V* | this study |
| VT4359 | *alg-2(ok304) II; lin-66(ok3326) IV; wIs51 V* | this study |
| VT4356 | *ain-2(tm1863) I; lin-66(ok3326) IV; wIs51 V* | this study |
| VT4497 | *ain-2(tm2432) I; lin-66(ok3326) IV; wIs51 V* | this study |
| VT4378 | *ain-2(tm1863) I; wIs51 V* | this study |
| VT4496 | *ain-2(tm2432) I; wIs51 V* | this study |
| VT4354 | *wIs51 V; ain-1(ku322) X* | this study |
| VT3751 | *maIs105 V; hbl-1(ma430[hbl-1::linker::mScarlet-I]) X* | Ambros Lab |
| VT4353 | *lin-66(ok3326) IV; wIs51 V; ain-1(ku322) X* | this study |
| VT4257 | *lin-66(ok3326) IV; lin-14(cc2841)* | this study |
| VT4350 | *lin-66(ok3326) IV; wIs51 V; hbl-1(ma430 scarlet)x* | this study |
| PD1301 | *lin-14(cc2841[lin-14::GFP]) X* | CGC |
| VT4398 | *lin-28(ma426[gfp]); lin-66(ok3326) IV*<br>*va* | this study |
| VT4399 | *lin-28(ma426)* | this study |
| VT4289 | *lin-28(ma562) I; lin-66(ok3326) IV; lin-14(cc2841) X.* | this study |
| VT4498 | *lin-28(ma582)I;lin-66(ok3326) IV; wIs51* | this study |
| VT4485 | *lin-28(ma582)I;wIs51* | this study |
| VT4232 | *lin-28(ma562)I;lin-66(ok3326) IV; wIs51* | this study |
| VT4484 | *lin-28(ma562); wIs51* | this study |
| VT4499 | *lin-28(n719) I;lin-66(ma646)IV;lin-46(ma385)V;*<br>*wIs51* | this study |
| VT4500 | *lin-66(ma646)IV;lin-46(ma385)V; wIs51* | this study |
| VT4501 | *lin-28(n719) I;lin-66(ma646)IV; wIs51* | this study |
| VT3818 | *lin-28(n719) I; lin-46(ma385) wIs51 V.* | Ambros Lab |
| VT3747 | *lin-46(ma385) wIs51 V.* | Ambros Lab |
| VT3442 | *lin-28(n719) I;wIs51 V.* | Ambros Lab |
| VT4502 | *wIs51 V: lin-14(n355n679) X* | this study |
| VT4478 | *lin-66(ok3326) IV; wIs51 V: lin-14(n355n679) X* | this study |
| VT4503 | *lin-28(ma562::GFP); lin-66(ok3326) IV; wIs51 V:*<br>*lin-14(n355n679) X* | this study |
| VT4504 | *lin-66(ok3326) IV; wIs51 V: lin-14(n355n679) X* | this study |
| VT4505 | *lin-28(ma416ma562); lin-66(ok3326) IV; wIs51 V* | this study |
| VT4506 | *lin-28(ma416ma562); wIs51 V: lin-14(n355n679)*<br>*X* | this study |
| VT3819 | *lin-28(n719); daf-7(e1372); lin-46(ma385); wIs51* | Ambros Lab |
| VT3746 | *daf-7(e1372); lin-46(ma385); wIs51* | Ambros Lab |
| VT3704 | *daf-7(e1372); wIs51* | Ambros Lab |
| VT4507 | *lin-28(ma416ma562); lin-66(ok3326) IV: lin-*<br>*14(n355n679) X* | this study |
| VT4508 | *lin-28(ma416ma562); lin-66(ok3326) IV* | this study |
| VT4509 | *lin-28(ma416ma562); lin-14(n355n679) X* | this study |

**Table S2:**
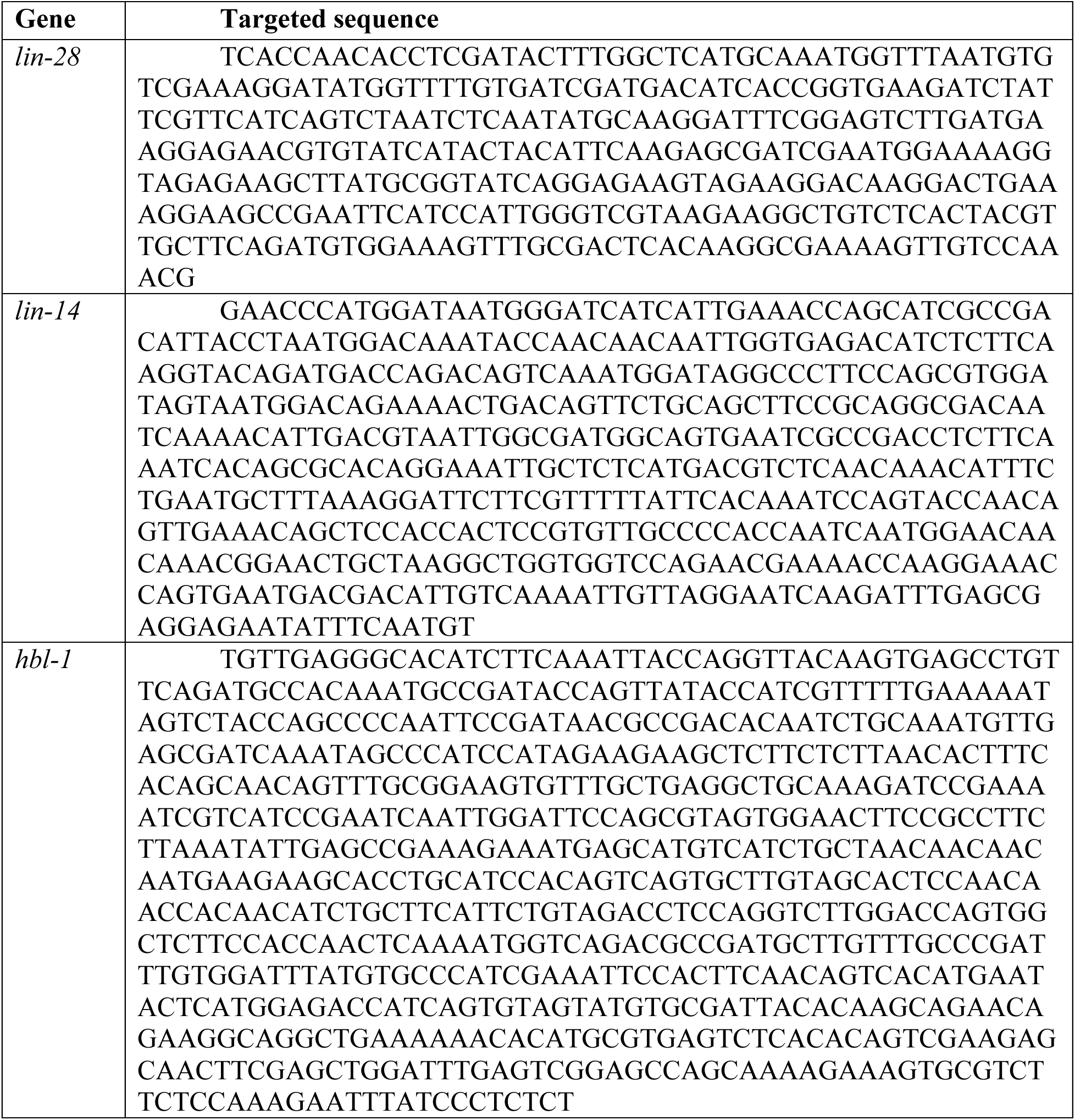
Targeted sequences in the newly designed Super RNAi vectors.

**Table S3:**
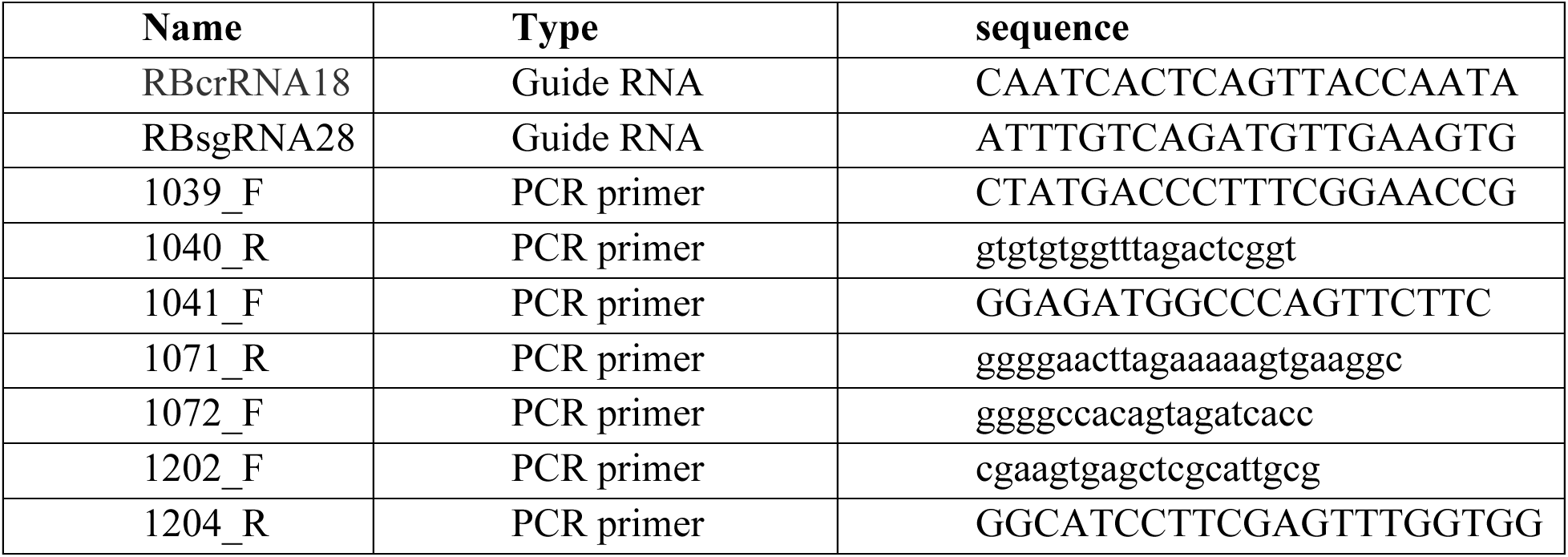
Oligos used in this study.

## Notes

### Competing Interest Statement

The authors have declared no competing interest.

